# An inflection point in high-throughput proteomics with Orbitrap Astral: analysis of biofluids, cells, and tissues

**DOI:** 10.1101/2024.04.26.591396

**Authors:** Nathan G. Hendricks, Santosh D. Bhosale, Angel J. Keoseyan, Josselin Ortiz, Aleksandr Stotland, Saeed Seyedmohammad, Chi D. L. Nguyen, Jonathan Bui, Annie Moradian, Susan M. Mockus, Jennifer E Van Eyk

## Abstract

This technical note presents a comprehensive proteomics workflow for the new combination of Orbitrap and Astral mass analyzers across biofluids, cells, and tissues. Central to our workflow is the integration of Adaptive Focused Acoustics (AFA) technology for cells and tissue lysis, to ensure robust and reproducible sample preparation in a high-throughput manner. Furthermore, we automated the detergent-compatible single-pot, solid-phase-enhanced sample Preparation (SP3) method for protein digestion, a technique that streamlines the process by combining purification and digestion steps, thereby reducing sample loss and improving efficiency. The synergy of these advanced methodologies facilitates a robust and high-throughput approach for cells and tissue analysis, an important consideration in translational research. This work disseminates our platform workflow, analyzes the effectiveness, demonstrates reproducibility of the results, and highlights the potential of these technologies in biomarker discovery and disease pathology. For cells and tissues (heart, liver, lung, and intestine) proteomics analysis by data-independent acquisition mode, identifications exceeding 10,000 proteins can be achieved with a 24-minute active gradient. In 200ng injections of HeLa digest across multiple gradients, an average of more than 80% of proteins have a CV less than 20%, and a 45-minute run covers ∼90% of the expressed proteome. In plasma samples including naive, depleted, perchloric acid precipitated, and Seer nanoparticle captured, all with a 24-minute gradient length, we identified 87, 108, 96 and 137 out of 216 FDA approved circulating protein biomarkers, respectively. This complete workflow allows for large swaths of the proteome to be identified and is compatible across diverse sample types.

**Graphical abstract created with biorender.com:** 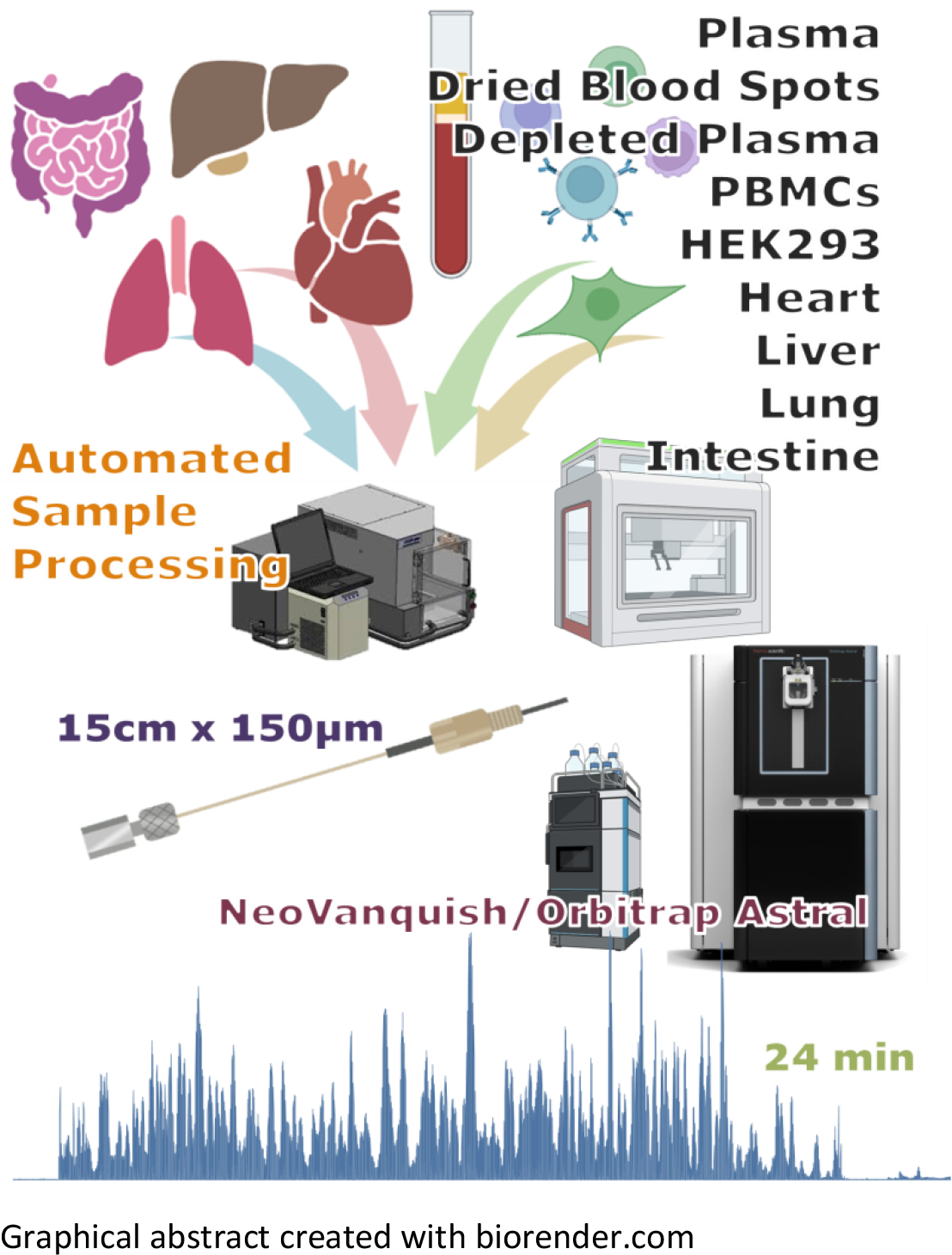

## Introduction

The introduction of the Orbitrap™ Astral™ mass spectrometer (MS) is a major technological feat in mass spectrometry due to a significant boost in the depth and throughput capabilities of LCMS-based proteomic workflows. Providing both high sensitivity, scan speed up to 200 Hz, and highly parallelizable acquisition, various groups have described impressive gains in protein identifications and throughput with good reproducibility.^1–11^ Users have reported remarkable levels of depth achievable through this new instrumentation in cellular, plasma and metaproteomic studies. In the context of clinical proteomics, large scale analysis of patient cohorts must combine precision, reproducibility, and robustness. Accordingly, we focused on leveraging the power of this new technology together with automated methods for sample lysis and digestion. Specifically, we describe methods and identify benchmarks for biospecimens in three categories, 1) Biofluids - plasma (naïve, depleted, perchloric acid-precipitated, and Seer nanoparticle-fractionated), dried blood spots, 2) Cells - HeLa, PBMCs, HEK293 and 3) mouse tissues – heart, intestine, liver, and lung.

To facilitate high-throughput analysis of these biospecimens, we have optimized the analysis conditions for a 15cm by 150μm ID Evosep (PepSep) column with 1.9μm particle size. To our knowledge, the use of this column has not been described with Vanquish LC and Astral MS. Current publications on the Orbitrap Astral MS primarily describe workflows with EasySpray (Pepmap) columns and standard methods for the NeoVanquish system.^11^ We have found that a Evosep column-oriented frontend to the Orbitrap Astral MS also offers reliable chromatography, run time utilization, and robustness.

## Experimental

### Biospecimens collections

Animal procedures were approved by the Cedars-Sinai Medical Center Institution Animal Care and Use protocol. Heart, intestine, liver, and lung samples were dissected from 129.S1 mice (Jackson Laboratory, Stock 002448). Frozen tissue was cut while on dry ice using blade or a disposable biopsy puncher (Integra Miltex Disposable Biopsy Puncher with plunger 1.5 mm), and ∼3-6 mg was used for lysis. Healthy human pooled plasma (BioIVT) was used with and without enrichment of low abundant proteins for proteomics analysis. HEK293 cells were procured from ATCC and maintained in DMEM supplanted with 10% FBS and antibiotic/antimycotic, in 5%CO2 incubator at 37C. Peripheral Blood Mononuclear Cells (PBMC) was acquired from Stem Cell Technologies and Pierce Hela protein digest standard from Thermo Fisher Scientific.

Various cell and tissue types were lysed using AFA technology on a Covaris LE220-plus sonicator (Covaris) in with either 8 AFA-tube TPX strips or m-130 glass tubes with beads depending on sample type. When working with less than one million cells, an 8 AFA-tube TPX strip (Covaris) was used. Cells were resuspended in 40 μL 100mM Tris-HCl pH 8 and 2% SDS in the TPX strip. After one minute centrifugation, the samples were sonicated using parameters set A values on the Covaris instrument (Supplementary Information). The total procedure time is 5 min per column. Afterwards, sample strips were centrifuged for 1 minute, then the supernatant was collected in a new tube and centrifuged for 20 min at max speed. For tissues and cell samples over one million cells, m-130 glass tubes with beads (Covaris) were used in place of the above procedure, but total volume was 120 μL and parameters set B values was used (Supplementary Information).

Cell and tissue lysates were digested by an automated SP3 protocol adapted to a Beckman i7 workstation. Bead aliquoting, reduction, alkylation, digestion, and elution were all performed on-deck with a 96-well plate format. Briefly, 20 μg of protein in 40 μL of the previously mentioned lysis buffer were reduced with the addition of 10 μL of 200 mM dithiothreitol and incubated 30 minutes at 60°C with shaking at 300 RPM, then alkylated with 10 μL of 400 mM IAA at room temperature for 15 minutes in the dark. The volume was brought to 70 μL with Tris HCl pH 8, then 5 μL of bead suspension (10:1 mass ratio of beads to protein, 1:1 mixture of hydrophilic/hydrophobic beads (Cytiva)) was aliquoted into the samples using the span-8 pipetting head, with constant agitation of the bead reservoir between transfers. The samples were brought to 50% ACN and incubated for 18 minutes, then solvent was removed on-magnet and samples were rinsed with 2x 80% EtOH then 2x ACN with 200 μL volumes each. After solvent was completely removed, samples were resuspended in 50 mM Tris HCl pH 8 and 10mM CaCl_2_ with trypsin at a 1:20 ratio. Samples were sonicated for 5 minutes then incubated 18 hours at 37°C and 1200 RPM overnight. After digestion, samples were then removed from beads and brought to 0.1% FA and 2% DMSO for injection onto the instrument.

Healthy pooled plasma samples were depleted using a 96-well filter plate (prototype plate, Thermo Fisher). 10 μL of pooled plasma was pipetted directly into the plate and 300 μL of Pierce Top-14 Depletion resin was added to the wells. Samples were incubated at room temperature with gentle mixing for 1 hour at 350 rpm. Samples were then eluted by positive pressure and dried down before digestion. Alternatively, samples were depleted by perchloric acid precipitation according to the protocol in Viode *et al*.^12^

Healthy pooled plasma samples were also prepared by treatment with proprietary nanoparticle from Seer Proteograph™, as described in Ferdosi *et al*.^8,13^ This automated platform offers a complete workflow from enrichment of low abundant proteins to peptide solution ready to be analyzed by LC-MS/MS using proprietary reagents.

5 μL of naïve plasma or dried depleted plasma samples were digested on the i7 automation workstation (Beckman Coulter). Samples were digested using the procedure described in Ardle *et al*.,^14^ using 1% SDC in place of TFE as a denaturing agent.

### LC-MS/MS analysis

Approximately 500 ng of digested samples were analyzed on a Thermo Orbitrap Astral coupled to a NeoVanquish LC. Samples were separated using 8-, 11-, 24-, 45-, or 60-minute gradients, with details given in supplementary information. The composition for solvent A and B were 0.1% FA in water and 80% ACN with 0.1% FA respectively. With all gradients the NeoVanquish LC was operated in direct injection mode, using a 150 μm ID x 15 cm 1.9 μm column (Evosep). The Evosep EasySpray adapter with a 30um ID metal emitter (Evosep) was coupled to a nano source (Thermo EasySpray) on the Orbitrap Astral MS platform (ThermoFisher).

All sample runs were acquired in data independent acquisition (DIA) mode from 380-980 Da with 240k Orbitrap resolution and 5ms maximum injection time for MS1. All DIA scans were set to 7ms maximum injection time with varying window schemes between 2 and 5 Th depending on gradient length.

### Data analysis

ProEpic™, an in-house platform that hosts a collection of proteomics workflows, was used to process MS raw data. The widely used proteomics search algorithms are inbuilt in the ProEpic™ v.2 environment.

Briefly, the MS raw data files were searched against UniProt human reviewed protein sequence entries (accessed April 2023) using DIA-NN (v 1.8.1)^15^ in library free mode with default parameters. Seer Proteograph™ (nanoparticle) samples were analyzed in Proteograph Analysis Suite, using a DIA-NN (v 1.8.1) environment in library-free mode. The output protein group matrix from DIA-NN was used to perform downstream analysis using Excel and R (see supplementary for additional details).

## Results and Discussion

For this study we assessed the depth and technical reproducibility of the NeoVanquish-Astral workflow with a 3x3 study design (three replicates on three separate days) of assorted tissue and biofluid sample types. We achieved optimal results using a 150 μm by 15 cm Evosep (Pepsep) column with 1.9 μm particle size in a direct-injection configuration. The moderate backpressure of this column enables fast loading even in direct injection and higher flow rates to elute peaks early and maximize the utilization of the gradient time (graphical abstract).

Table 1 illustrates the total identifications (as retrieved from the DIA-NN PG matrix) for all 9 replicates per sample type, gradient and MS method. HeLa samples were injected at 200ng—a typical benchmarking sample load used across many instruments for all gradients and methods. We focused on HeLa and plasma (depleted and naïve) for different run times and DIA method parameters to explore tradeoffs in coverage, reproducibility, and throughput. The high scan speed of the Astral mass analyzer translates to lower cycle times, enabling small DIA windows with short gradients while still sampling many points across the analyte elution peak. We explored the tradeoffs in pushing these limits while comparing two DIA method parameters in most gradients: one method with a quantitation-optimized window size (e.g. 24-minute 3Th) giving ≥6 points across the peak on average, and another with a smaller window size (e.g. 24-minute 2Th) and longer cycle time. In all cases, we see more identifications with a smaller window size, but we also examined the impact on quantitative accuracy calculated as % coefficient of variance (%CV). As seen in Figure 1A, the impact of window size for the 24-minute gradient noticeably skews the % CV distribution when comparing 2 versus 3 Th windows. Still, with the less quantitatively robust 2 Th window size the average %CV is below 13% for HeLa and 20% for Plasma. Interestingly, in shorter gradient times the impact of window size on %CV is less pronounced, particularly with the 11-minute gradient where there is little difference in the two window sizes, and in general shorter gradients have tighter CV distributions. For HeLa, a 45-minute gradient covers approximately 90% of the expressed proteome,^16^ while an 8-minute gradient can deliver 78% as many hits in less than 1/5 of the time. In terms of copy numbers per cell, this level of coverage can observe proteins at ∼100 copies, with the lowest observed protein at an estimated 13 copies per cell (supplementary figure 1). Furthermore, this deep coverage of the proteome uncovers some of the “missing proteins” in the human proteome—proteins that are expected to be present based on the transcriptome or genome but lack direct protein-level evidence.^17,18^ Five of these proteins were detected with two or more peptides >9 amino acids in length, meeting the stringent criteria to be moved to Protein Existence level 1 (supplementary Table 1).^19^

**Table 1:** The total protein group (PG) identifications in each sample type with respect to the different gradients and MS methods tested. Numbers are given for 9 total injections (3 replicates by 3 days), except for the Seer nanoparticle runs, where 3 replicates of five-nanoparticle fractions were analyzed on the given gradient for a total of 15 injections on one day.

| Gradient<br>(min) | Window | HeLa<br>(200ng) | Naïve<br>Plasma | Depleted<br>Plasma | PerCA<br>Plasma | Seer<br>nanoparticle | PBMCs | Dried Blood<br>Spots | HEK293<br>Cells | Mouse<br>Heart | Mouse Lung | Mouse Liver | Mouse<br>Intestine |
| --- | --- | --- | --- | --- | --- | --- | --- | --- | --- | --- | --- | --- | --- |
| 8 | 5 | 8126 | 475 | 834 |  |  |  |  |  |  |  |  |  |
| 8 | 4 | 8265 | 565 | 837 |  |  |  |  |  |  |  |  |  |
| 11 | 4 | 8598 | 572 | 974 |  |  |  |  |  |  |  |  |  |
| 11 | 3 | 8700 | 636 | 1051 |  |  |  |  |  |  |  |  |  |
| 24 | 3 | 9353 | 695 | 1198 | 1567 | 3359* | 8157 | 2693 | 10140 | 7056 | 10808 | 6758 | 8879 |
| 24 | 2 | 9817 | 896 | 1307 |  |  |  |  |  |  |  |  |  |
| 45 | 2 | 10619 | 1177 | 1541 |  |  |  |  |  |  |  |  |  |

**Figure 1:**
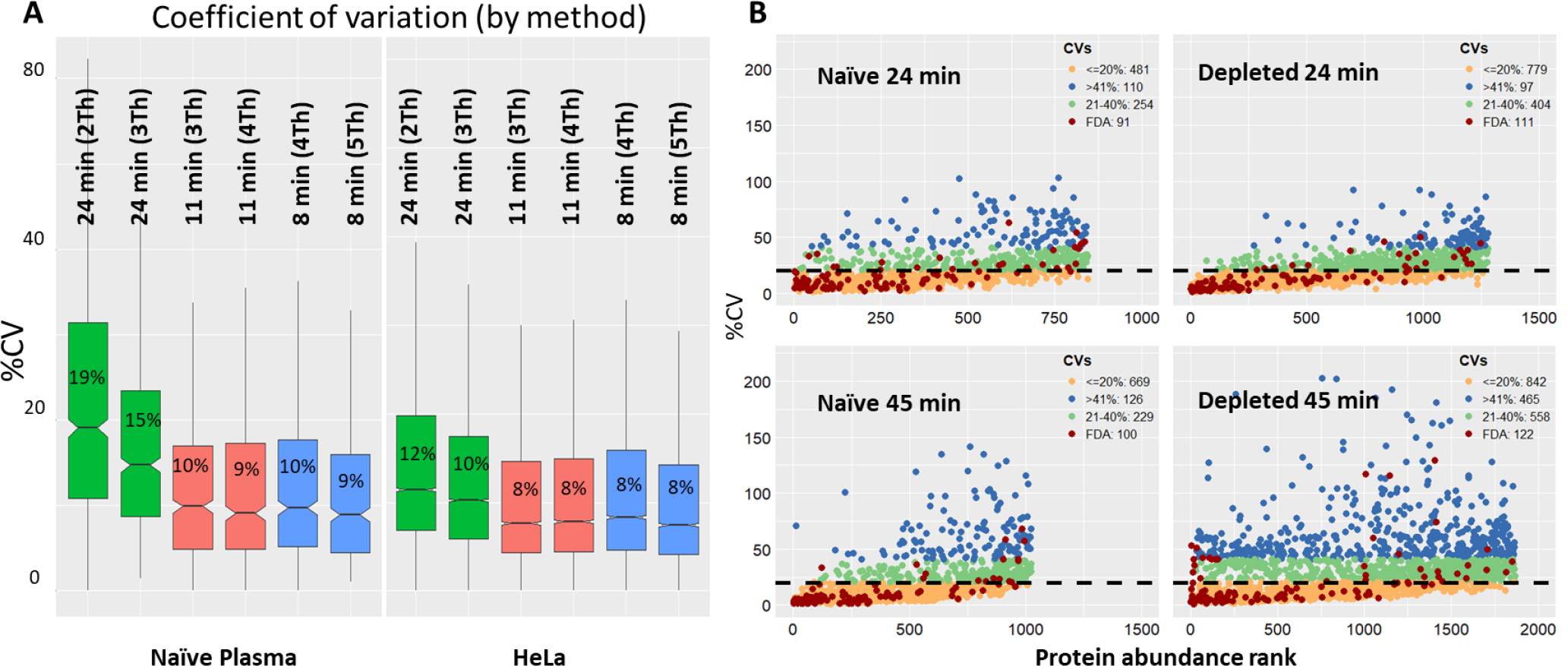
**A)** Comparison of the different gradients and DIA window sizes by %CV. Box plots are given for naive plasma (left) and HeLa (right) according to three different gradients, with two window sizes per gradient. Median %CVs are overlaid. **B)** CV plots of all identification in naïve (right) and depleted plasma (left), in both 24 (top) and 45-minute (bottom) runs. Published FDA biomarkers are listed highlighted in red.

The high dynamic range of plasma still poses difficulty for deep proteomic analysis, but protein identifications in naïve plasma still represent an impressive technical advancement, with over 1100 proteins seen in a 45-minute gradient. Depletion offers considerable gains in depth particularly in shorter runs, giving more than 350 additional identifications in an 8-minute gradient. Impressively, the perchloric acid (PerCA) precipitation approach to depletion delivers over 1500 identifications. Additionally, Seer nanoparticles were run on pooled plasma and analyzed with the 24-minute gradient. Three replicates were analyzed, but each replicate represents 5 injections (for each of the nanoparticles). These different approaches to increasing the depth of coverage in plasma are compared alongside Naïve with 24-minute gradient (Figure 2A). The preparations we explored are evidently complimentary, with no single method having total overlap with the others (Figure 2B).

**Figure 2:**
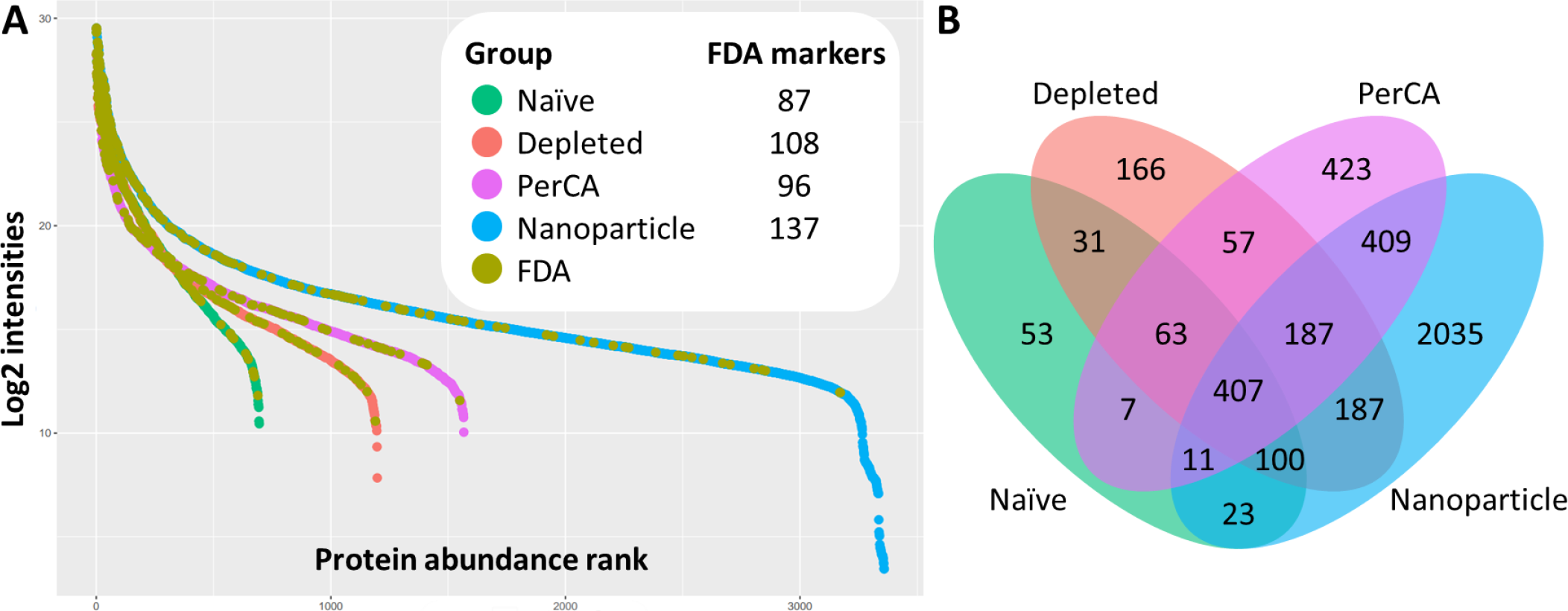
**A)** Protein abundance rank plots of the of naïve and depleted plasma samples in the 24-minute 3Th MS method. **B)** Proteins seen in antibody-depleted of top 14 abundant plasma proteins, PerCA precipitated, and Seer Proteograph™ plasma samples and their overlap with Naïve are shown in a Venn diagram.

We also investigate the coverage of known biomarkers in plasma that are currently used in clinical chemistry laboratories for clinical decision making. Figure 1B shows FDA-approved circulating biomarkers^20^ as seen in naïve and depleted plasma in 24- and 45-minute gradients arranged by %CV and protein rank according to the list of 216 described in Bhowmick *et al*. Between 91 and 122 biomarkers are seen per sample set, the majority of which lie below 20% CV. Even in healthy pooled plasma, low-abundance markers such as ovarian carcinoma antigen CA125 (Mucin-16, uniprot: Q8WXI7), Pancreatic triacylglycerol lipase (PNLIP, uniprot: P16233), and Platelet glycoprotein 4 (CD36, uniprot: P16671) are observed. In addition to biofluids, tissue and cell samples hold significant importance in the development and discovery of biomarkers in biopsy samples and fundamental research of diagnostics and therapeutics. We analyzed an assortment of different tissue and cell types with the 24-minute 3 Th gradient. Heart, liver, lung, and intestine samples from mice were processed in an automated, high throughput sample preparation workflow. AFA sonication technology, capable of bulk sonication of samples in plate format, was used for efficient sample lysis of tissues in 2% SDS. Sample lysates were then digested by automated SP3^12,21,22^ on the Beckman i7 automated workstation. The Beckman i7 has 96 and 8-channel pipettor heads, along with on-deck shaking, incubation, and magnetic pulldown of SP3 beads. The protocol we developed for this study includes use of the span-8 pipettor and orbital shaker to perform bead aliquoting into sample wells on-deck, which has not been previously described to our knowledge, removing all the laborious bead-handling steps associated with SP3 (SI figure 2).

PBMC’s consist of two main types of white blood cells lymphocytes (including B, T and NK cells) and monocytes. Studying PBMCs provides a detailed understanding of how specific cells respond to various stimuli such as infections, cancers, or therapeutic drugs.^23,24^ To the best of our knowledge, close to 9000 protein identification from PBMCs represents the most extensive proteome coverage achieved in a 24-minute analysis. Figure 3 shows highly significant enrichment of reactome pathways seen in PBMCs covering metabolism, disease, cell cycle, and immune response, reflecting the variety of responses to stimuli that could be interrogated in a 24-minute analysis.

**Figure 3:**
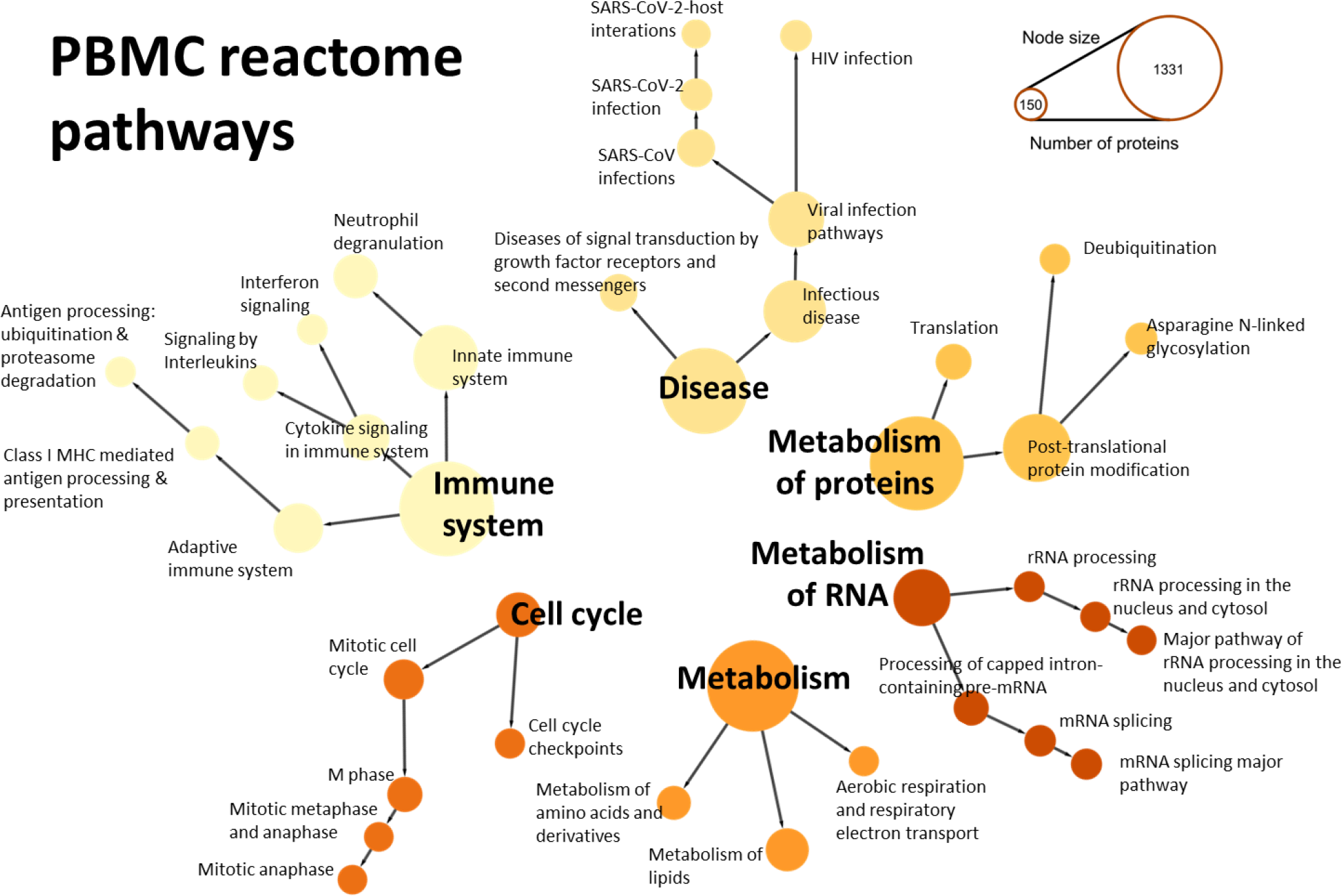
reactome pathways based on the molecular function of proteins identified in PBMC samples on the Astral 24-minute runs. Node size represents number of proteins and is filtered to display only pathways with 150 proteins or more. All nodes have significant adjusted p-value enrichment, with a range of 3.6e-3 to 9.9e-141.

Also of note is the number of identifications in heart tissue, which is a typically challenging for analysis due to the high-abundant sarcomeric myofilament proteins responsible for heart contractile motion that obscure low-abundant species, acting much like albumin in plasma.^25,26^ When searched together with the other murine tissues, we identify over 10k protein groups corresponding to 8928 unique proteins. The ontology of these identifications is displayed in figure 4, with coverage of the Golgi apparatus, nucleus and mitochondria inset. More than 800 of the estimated 1100-1400 mitochondrial proteins are identified in heart tissue.^27^

**Figure 4:**
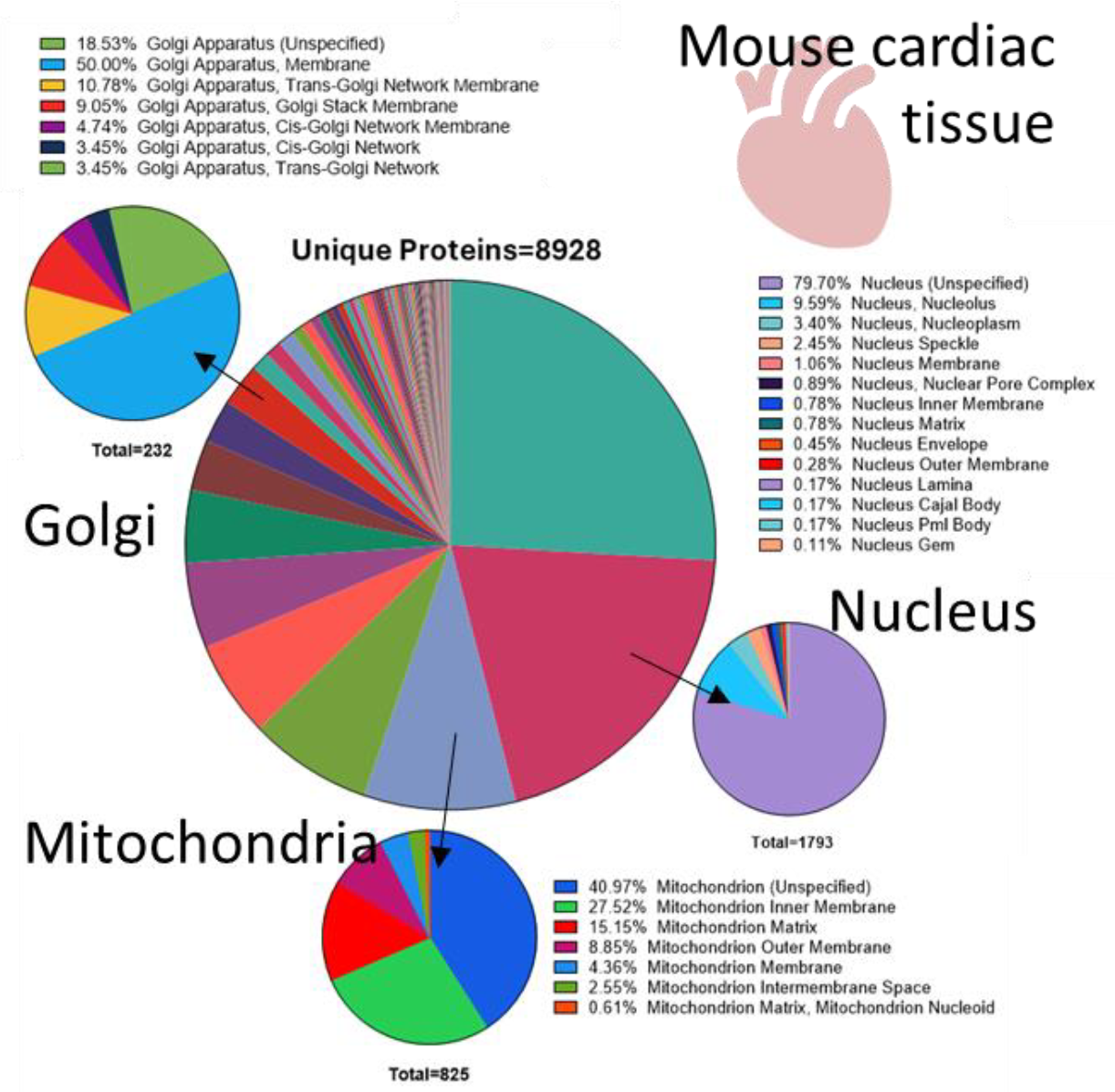
Gene ontology of the identifications in murine heart tissue samples when co-analyzed with other murine tissues in DIA-NN. Coverage of Golgi, mitochondrial, and nuclear proteins within each section are further expanded.

## Conclusions

Our study outlines efficient, high-throughput workflows for automated sample preparation and rapid analysis of diverse biospecimens using the Orbitrap Astral MS platform. A significant breakthrough lies in our ability to characterize approximately 90% of a sample proteome of cells and tissue within a mere 24 -minute gradient. Our fast proteomics analysis of biofluids such as whole blood, naive, and depleted plasma demonstrates high-quality quantitative measurements across a wide dynamic range. Notably, our results demonstrate that CVs perform exceptionally well at short gradients, enabling high-throughput analyses with impressive precision. The remarkable sensitivity and rapid scan speed of the Orbitrap Astral reveal previously unexplored depths, allowing robust identification and quantification of low-abundance proteins. These capabilities open new opportunities in biomarker discovery and drug development. In summary, the study’s findings pave the way for more efficient and comprehensive proteomic analyses across diverse sample types, with potential implications for advancing medical research and therapeutic innovations.

## Acknowledgments

We acknowledge Lior Zilberberg for providing mouse tissues, Christopher Hughes, Sandi Spencer Miko and Gregg Morin, for insightful discussions on troubleshooting the SP3 protocol, Tony Herren for providing Thermo’s prototype filter plates for plasma depletion and Sameer Vasantgadkar for all the help with Covaris instrumentation setup, discussions and suggestions for protocol improvements of cell lysis procedure on Covaris.

## Funding

NIH R01HL111362 (JVE), U01 DK124019-01 (JVE) and R01 HL155346-01A1 (JVE), Erika Glazer Endowed Chair (JVE), and Cedars-Sinai Academic Affairs, Proteomics and Metabolomic core and Precision Health.

## Supplementary Information

### Covaris Instrument Parameters

**SI figure 1:**
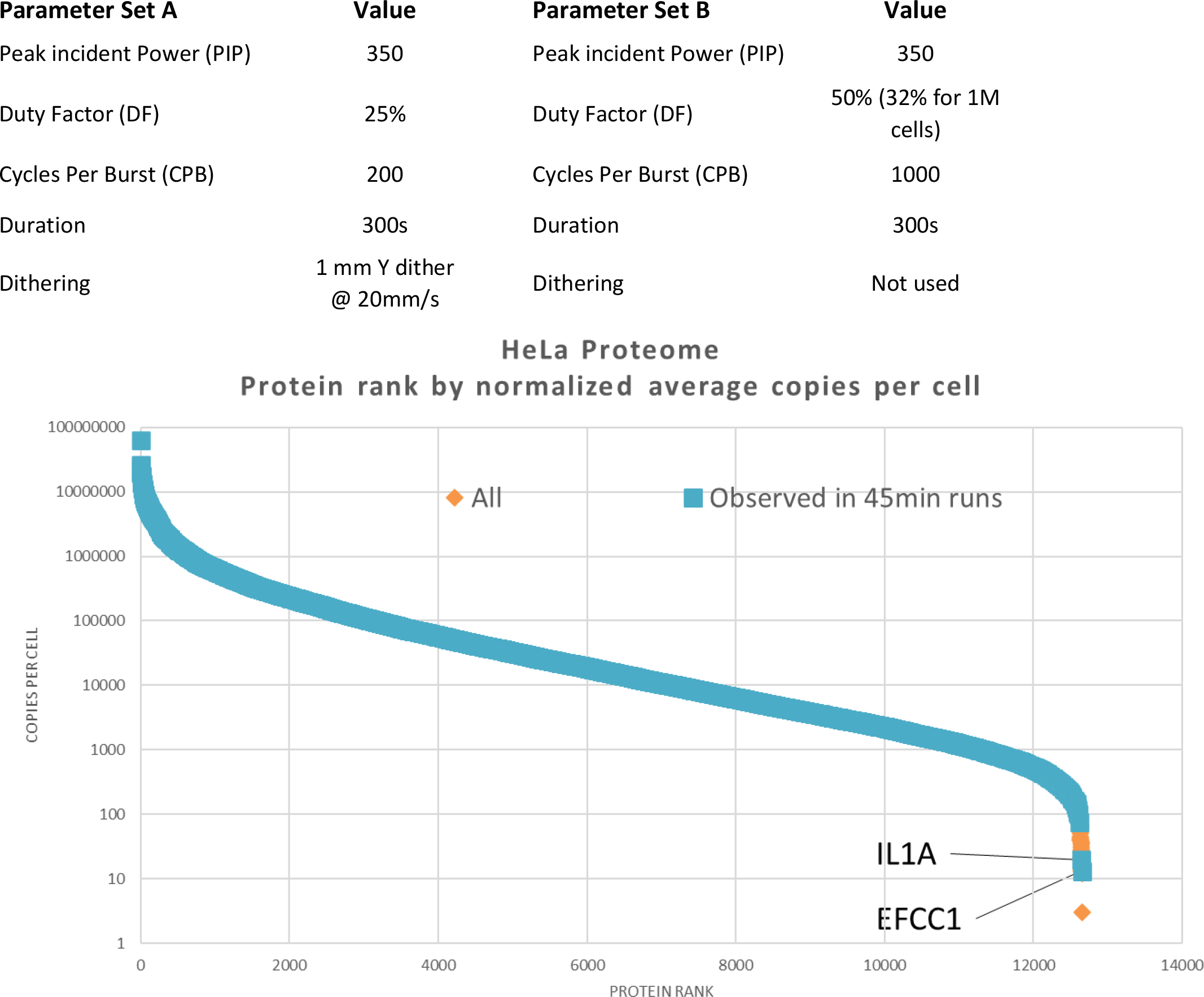
dynamic range coverage of the total HeLa proteome in according to estimated copies per cell, as seen in 200ng on a 45-minute gradient. 10,619 of the total 12,654 proteins described in Dolgalev *et al*. are seen (blue markers). The least abundant identified protein is EFCC1 at 13 copies per cell.

**SI table 1:**
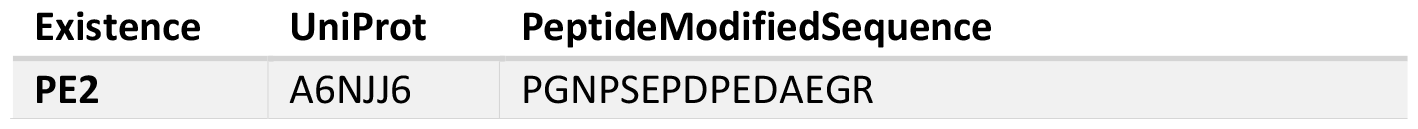
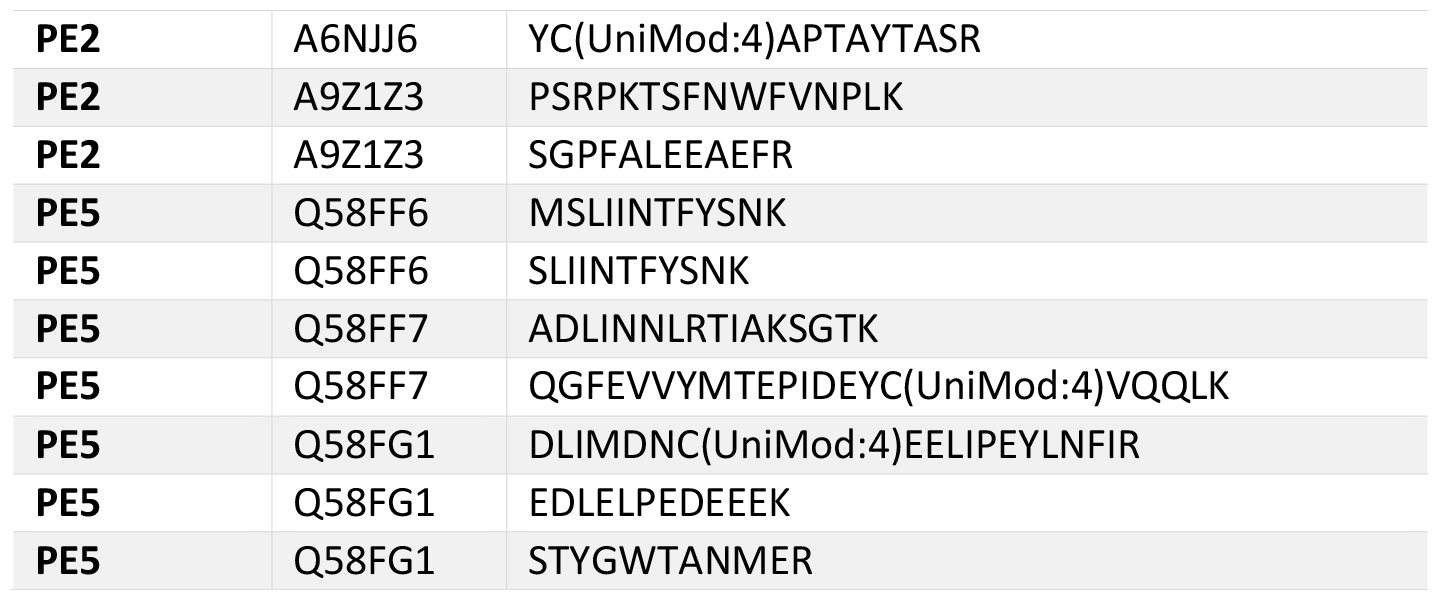
“Missing” human proteins observed in the search of HeLa data, including peptide sequences with more than 9 amino acids.

**SI Figure 2:**
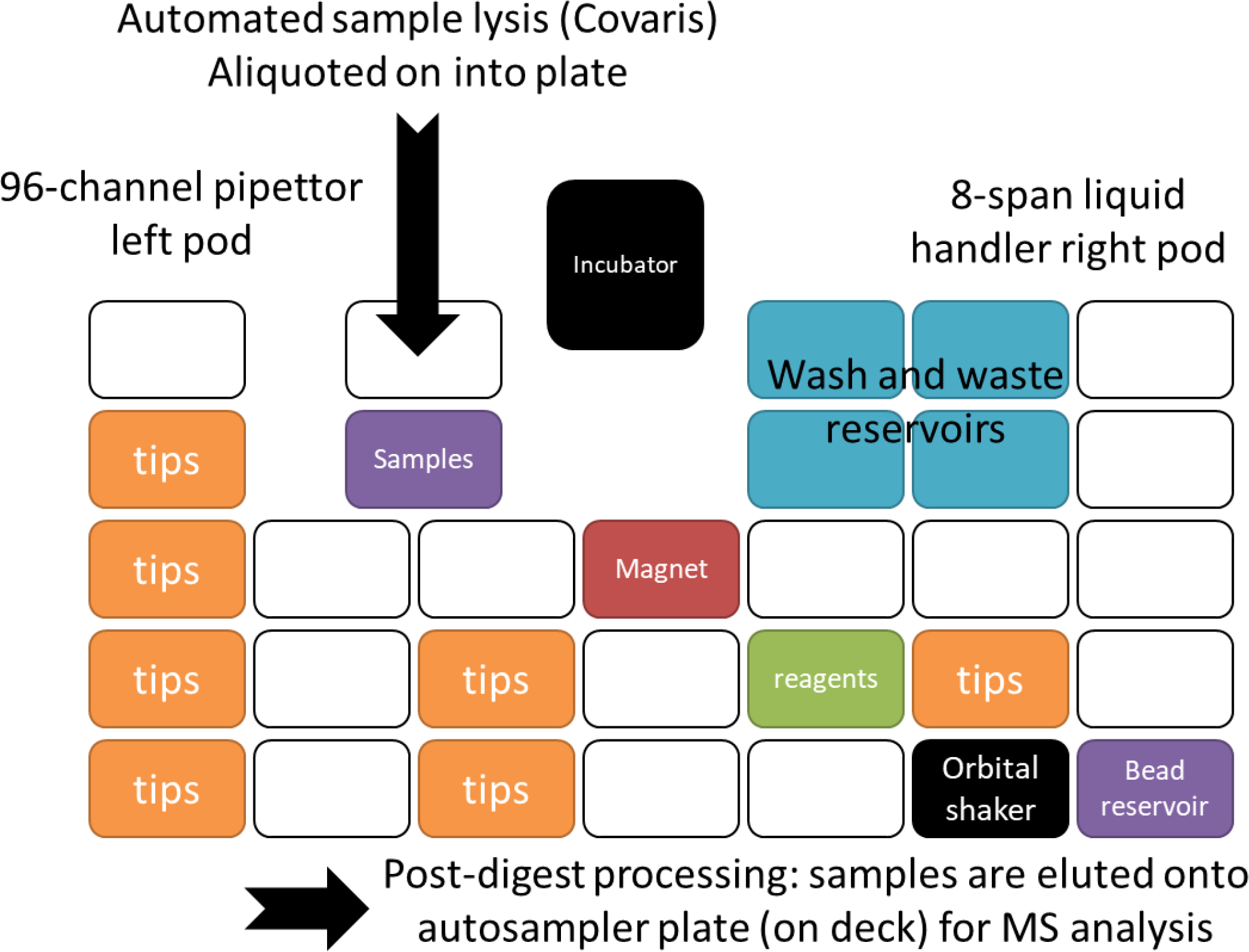
Starting layout of Biomek i7 deck. Incubation, shaking, bead aliquoting, bead rinsing, and bead removal all occur on-deck.

## Data analysis of PBMCs for pathway enrichment

Gene set analysis (GSA) using the piano R package version 2.18.0 developed by Väremo and colleagues was applied in this study. The runGSAhyper function was applied for the statistical testing of the overrepresented pathways using Fisher’s exact test to find enriched pathways. This function returned the protein set P-values indicating the significance enrichment of a pathway. Reactome pathways database was used for the pathway enrichment analysis. The entire mapped pathways including 6245 proteins mapped to 2679 pathways were used as background information. Only pathways with a minimum of 150 proteins and an adjusted p value (Benjamin-Hochberg correction) of less than 0.005 were considered in further analysis.

All the edges represent the parent-child pathway relationship. The pathway illustration was performed using Cytoscape version 3.9.1 (Shannon et al. 2003) via RCy3 R package version 2.22.1 (Gustavsen et al. 2019). The hierarchical layout was applied for parent-child pathway interaction.

## Notes

### Competing Interest Statement

The authors have declared no competing interest.

## References

1. Stewart, H. I. et al. Parallelized Acquisition of Orbitrap and Astral Analyzers Enables High-Throughput Quantitative Analysis. Anal. Chem. 95, 15656–15664 (2023).

2. Grinfeld, D. et al. Multi-reflection Astral mass spectrometer with isochronous drift in elongated ion mirrors. Nucl. Instrum. Methods Phys. Res. Sect. Accel. Spectrometers Detect. Assoc. Equip. 1060, 169017 (2024).

3. Bortel, P. et al. Systematic optimization of automated phosphopeptide enrichment for high-sensitivity phosphoproteomics. Mol. Cell. Proteomics 100754 (2024) doi:10.1016/j.mcpro.2024.100754.

4. Stewart, H. et al. A Conjoined Rectilinear Collision Cell and Pulsed Extraction Ion Trap with Auxiliary DC Electrodes. J. Am. Soc. Mass Spectrom. 35, 74–81 (2024).

5. Yang, X., Zheng, X., Zhai, X., Tang, T. & Yu, S. Spindle apparatus coiled-coil protein 1 (SPDL1) serves as a novel prognostic biomarker in triple-negative breast cancer. PROTEOMICS – Clin. Appl. n/a, 2300002.

6. Heil, L. R. et al. Evaluating the Performance of the Astral Mass Analyzer for Quantitative Proteomics Using Data-Independent Acquisition. J. Proteome Res. 22, 3290–3300 (2023).

7. Dumas, T. et al. The astounding exhaustiveness and speed of the Astral mass analyzer for highly complex samples is a quantum leap in the functional analysis of microbiomes. Microbiome 12, 46 (2024).

8. Huang, T. et al. Protein Coronas on Functionalized Nanoparticles Enable Quantitative and Precise Large-Scale Deep Plasma Proteomics. 2023.08.28.555225 Preprint at 10.1101/2023.08.28.555225 (2023).

9. Serrano, L. R. et al. The one hour human proteome. Mol. Cell. Proteomics 100760 (2024) doi:10.1016/j.mcpro.2024.100760.

10. Ivanov, M. V., Kopeykina, A. S. & Gorshkov, M. V. Reanalysis of Orbitrap Astral DIA data demonstrates the capabilities of MS/MS-free proteomics to reveal new biological insights in disease-related samples. 2024.04.01.587550 Preprint at 10.1101/2024.04.01.587550 (2024).

11. Guzman, U. H. et al. Ultra-fast label-free quantification and comprehensive proteome coverage with narrow-window data-independent acquisition. Nat. Biotechnol. 1–12 (2024) doi:10.1038/s41587-023-02099-7.

12. Viode, A. et al. A simple, time- and cost-effective, high-throughput depletion strategy for deep plasma proteomics. Sci. Adv. 9, eadf9717 (2023).

13. Ferdosi, S. et al. Engineered nanoparticles enable deep proteomics studies at scale by leveraging tunable nano–bio interactions. Proc. Natl. Acad. Sci. 119, e2106053119 (2022).

14. Mc Ardle, A. et al. Standardized Workflow for Precise Mid- and High-Throughput Proteomics of Blood Biofluids. Clin. Chem. 68, 450–460 (2022).

15. Demichev, V., Messner, C. B., Vernardis, S. I., Lilley, K. S. & Ralser, M. DIA-NN: Neural networks and interference correction enable deep proteome coverage in high throughput. Nat. Methods 17, 41–44 (2020).

16. Dolgalev, G. V., Safonov, T. A., Arzumanian, V. A., Kiseleva, O. I. & Poverennaya, E. V. Estimating Total Quantitative Protein Content in Escherichia coli, Saccharomyces cerevisiae, and HeLa Cells. Int. J. Mol. Sci. 24, 2081 (2023).

17. Adhikari, S. et al. A high-stringency blueprint of the human proteome. Nat. Commun. 11, 5301 (2020).

18. Omenn, G. S. et al. The 2023 Report on the Proteome from the HUPO Human Proteome Project. J. Proteome Res. 23, 532–549 (2024).

19. Deutsch, E. W. et al. Human Proteome Project Mass Spectrometry Data Interpretation Guidelines 3.0. J. Proteome Res. 18, 4108–4116 (2019).

20. Bhowmick, P., Roome, S., Borchers, C. H., Goodlett, D. R. & Mohammed, Y. An Update on MRMAssayDB: A Comprehensive Resource for Targeted Proteomics Assays in the Community. J. Proteome Res. 20, 2105–2115 (2021).

21. Hughes, C. S. et al. Single-pot, solid-phase-enhanced sample preparation for proteomics experiments. Nat. Protoc. 14, 68–85 (2019).

22. Müller, T. et al. Automated sample preparation with SP3 for low-input clinical proteomics. Mol. Syst. Biol. 16, e9111 (2020).

23. Prinz, C. Importance of Peripheral Blood Mononuclear Cells (PBMCs) in Clinical Immunology. J. Clin. Chem. Lab. Med. 5, 1–2 (2023).

24. Yang, L., Weng, S., Qian, X., Wang, M. & Ying, W. Strategy for Microscale Extraction and Proteome Profiling of Peripheral Blood Mononuclear Cells. Anal. Chem. 94, 8827–8832 (2022).

25. Karpov, O. A. et al. Proteomics of the heart. Physiol. Rev. 104, 931–982 (2024).

26. Ai, L. et al. High-Field Asymmetric Waveform Ion Mobility Spectrometry: Practical Alternative for Cardiac Proteome Sample Processing. J. Proteome Res. 22, 2124–2130 (2023).

27. Baker, Z. N., Forny, P. & Pagliarini, D. J. Mitochondrial proteome research: the road ahead. Nat. Rev. Mol. Cell Biol. 25, 65–82 (2024).

